# Tryptophan-like side chain holding aptamers inhibit respiratory syncytial virus infection of lung epithelial cells

**DOI:** 10.1101/2022.12.09.519757

**Authors:** Krisztina Percze, Zoltán János Tolnai, Marc Eleveld, Li Ou, Haijuan Du, Adam S. Olia, Peter D. Kwong, Marien I. de Jonge, Tamás Mészáros

## Abstract

Respiratory syncytial virus (RSV) is a leading cause of serious and even fatal acute lower respiratory tract infections in infants and in the elderly. Potent RSV neutralization has been achieved by antibodies that selectively bind the prefusion form of the viral fusion (F) protein. We hypothesised that similar potent neutralization could be achieved using F protein targeting aptamers. Aptamers have yet to reach their translational potential for therapeutics or diagnostics due to their short half-life and limited range of target-aptamer interactions; these shortcomings can, however, be ameliorated by application of amino acid-like side chain holding nucleotides. In this study, a stabilized version of the prefusion RSV F protein was targeted by aptamer selection using an oligonucleotide library holding a tryptophan-like side chain. This process resulted in aptamers that bound the F protein with high affinity and differentiated between pre- and postfusion conformation. Identified aptamers inhibited viral infection of lung epithelial cells. Moreover, introduction of modified nucleotides extended aptamer half-lives. Our results suggest that targeting aptamers to the surface of viruses could yield effective drug candidates, which could keep up with the pace of the continuously evolving pathogens.

## 1. INTRODUCTION

Respiratory diseases are a leading cause of morbidity and mortality among young children. Respiratory syncytial virus (RSV) is the most common seasonal respiratory virus usually leading to mild, cold-like symptoms in infants (1); however, it can also cause severe respiratory tract diseases, in particular bronchiolitis, inflammation of the small airways in the lungs and pneumonia (2). Besides young children, RSV infection can result in severe disease in immunocompromised hosts and the elderly and has been associated with the development of asthma (3). RSV infections are the leading cause of hospitalization among young infants worldwide and the second most common cause of child mortality in low- and middle-income countries (4).

RSV is a highly contagious pathogen and spreads among groups of young children, within families and between hospitalized patients via direct physical contact, droplets and aerosol transmission (5,6). Recent restrictive measures due to the COVID-19 pandemic have contributed to the disruption of the seasonal infection pattern of respiratory viruses. This may have led to an increase in the immunologically naïve population, causing more severe RSV outbreaks worldwide (7). Currently there is no licensed vaccine available to prevent RSV infections, while many vaccine candidates are under development. The only approved prophylaxes are monoclonal antibodies directed against F protein.

RSV is a pleomorphic enveloped RNA virus. Out of the 11 RSV genome encoded proteins, three (F, G, and SH) are anchored in the membrane and involved in virus entry. The two major surface exposed glycoproteins G and F are crucial for virus attachment and fusion, respectively. The small hydrophobic (SH) protein is suggested to be involved in ion channel formation (viroporin) to modulate the infected cell; however, the exact function is not entirely understood (8). The F protein is indispensable for infection of the host cell and its genetic diversity is low compared to protein G (9). These features make the F protein a rational target for therapeutic intervention. Conformation of protein F undergoes significant changes during the fusion of the membranes, i.e., the prefusion form of the protein is converted into post-fusion F. This process can also take place spontaneously indicating the metastability of the prefusion conformation (10). A multitude of human antibodies was raised against protein F and the prefusion F specific antigenic site Ø binding variants have been proven to possess strong virus neutralizing capability (11). To aid vaccine development, a stabilized version of prefusion protein F was created by structure-based design (12). This mutant version of RSV F maintained availability of the antigenic site Ø even when exposed to extremes of pH, osmolality, and temperature.

More than 200 viruses are capable of infecting humans and many of them are known etiologic agents of various diseases (13). Due to the high mutation rate of some viruses, there is a strong demand for developing novel drugs that can effectively block cell invasion or replication of the constantly emerging novel virus strains. Although the main target of RSV vaccine development, the F protein is characterized by a relatively high genetic stability, amino acid substitution that results in the emergence of prophylaxis resistant virus strains has also been described (14). The presently available antiviral drugs are dominated by small molecules which specifically bind to the surface proteins or replication machinery of viruses (15). Concerning macromolecular therapeutics, there is only a single oligonucleotide-based antiviral drug, which is used to treat cytomegalovirus infections and represent the first approved antisense therapy (16).

Aptamers are single-stranded oligonucleotides that can bind to their target molecules with similar affinity and specificity to those of antibodies. However, selection and synthesis of aptamers do not rely on living organisms thus providing advantages over antibodies in terms of time and cost of production (17). Virologists also realized the advantages of the relatively rapid and straightforward selection process of aptamers, the so-called SELEX, at the dawn of aptamer science. In fact, the very first published aptamers were generated by using the T4 bacteriophage DNA polymerase as a target molecule of selection (18). Since then, a wide array of human pathogen targeting aptamers has been described (19,20). Although the majority of published virus specific aptamers were made for diagnostic applications and applied in development of biosensor, there are also numerous examples for aptamers of antiviral potential (21). The first antiviral aptamer was raised against the reverse transcriptase of human immunodeficiency virus type 1 (HIV-1) to inhibit its replication (22). The reverse transcriptase specific RNA oligonucleotide was followed by selection of aptamers for a panel of human pathogen viruses. The target molecules of these SELEX process varied from recombinant viral surface and replication proteins to inactivated viruses and virus like nanoparticles and even to virus glycoprotein expressing mammalian cells (23–26).

The possible interactions between proteins and natural oligonucleotides are constrained due the limited chemical diversity of the nucleobases (27). Therefore, a series of unnatural modifications have been incorporated into aptamers to improve the efficiency of SELEX and increase the durability of aptamers in the prevailing environmental and physiological conditions. For modified aptamer selection, oligonucleotide libraries of nonstandard nucleotides have been created either by directly introducing the modified nucleotides during the synthesis or by the addition of “clickable” nucleotides and the post-synthetic functionalization by click-chemistry (27,28). According to the modification affected components of nucleotides, three broad categories, i.e., sugar, phosphodiester backbone, and nucleobase modified aptamers can be distinguished (29). Amongst the numerous applied unnatural oligonucleotides, the so-called SOMAmers (slow off-rate modified aptamers, base modified oligonucleotides at the 5-position of deoxyuridine (30)), have been proven to be the most promising candidates. Modified nucleotides of SOMAmers hold hydrophobic or aromatic functional groups at positions oriented away from the hydrogen bonding sites of the bases. Introduction of these modified nucleotides significantly increased the success rate of SELEX; moreover, most of the SOMAmers exhibit improved, sub-nanomolar K_D_ values (30).

Previously, we selected aptamers by using inactivated virus particles and demonstrated that the obtained aptamers specifically bind to protein G of RSV and can be applied for virus detection in clinically relevant samples (31). In this study, we set out to produce aptamers of therapeutic potential by applying a different selection rationale. The target molecule serving whole virus was replaced by a stabilized version of the prefusion protein F during the selection steps and the tryptophan-like side chain holding TAdUTP was introduced into the oligonucleotide library of SELEX. The selection process resulted in aptamers that could bind to their target protein with nanomolar dissociation rates; in addition, these aptamers could differentiate between the pre- and postfusion variant of protein F. Furthermore, the best aptamers were found to inhibit RSV infection with a similar effectivity to that of palivizumab.

## 2. RESULTS

### 2.1. Selection of aptamers

Considering that RSV infection can be prevented by prefusion F specific antigenic site Ø binding antibodies, we hypothesised that a similar effect could be achieved by prefusion F protein selective aptamers. To produce aptamers of therapeutic potential for RSV, we utilized a stabilized version of prefusion and the postfusion protein F as targets and counter-target of SELEX, respectively. To increase efficacy of the selection, we created a 5-indolyl-AA-dUTP (TAdUTP) possessing starting oligonucleotide library as detailed previously (32). The enriched oligonucleotide pools of seven SELEX cycles were amplified by emulsion PCR and by replacing dTTP with TAdUTP. Selection pressure was ensured by the addition of competitor molecules, e.g., L-tryptophan, mucin, BSA, Salmon DNA to the buffer and decreasing target-molecule concentration and incubation time in consecutive SELEX cycles. Furthermore, counter-selection steps were introduced by challenging the oligonucleotide pool with bead-immobilized postfusion F protein to favour aptamers of prefusion conformation selectivity (Figure 1). Following completion of the final selection step, nucleotide sequences of 96 aptamer candidates were determined. Computational analysis of the obtained sequences revealed that 10 aptamers were present in multiple copies, hinting enrichment of potential RSV selective modified aptamers. Although the MEME Motif search did not identify any consensus sequence, the modified nucleotide and guanosine were overrepresented in comparison to cytidine and adenine in all selected aptamer candidates (Figure S1). This finding suggests the necessity of the modified nucleotide in the aptamer-protein F complex formation.

**Figure 1.**
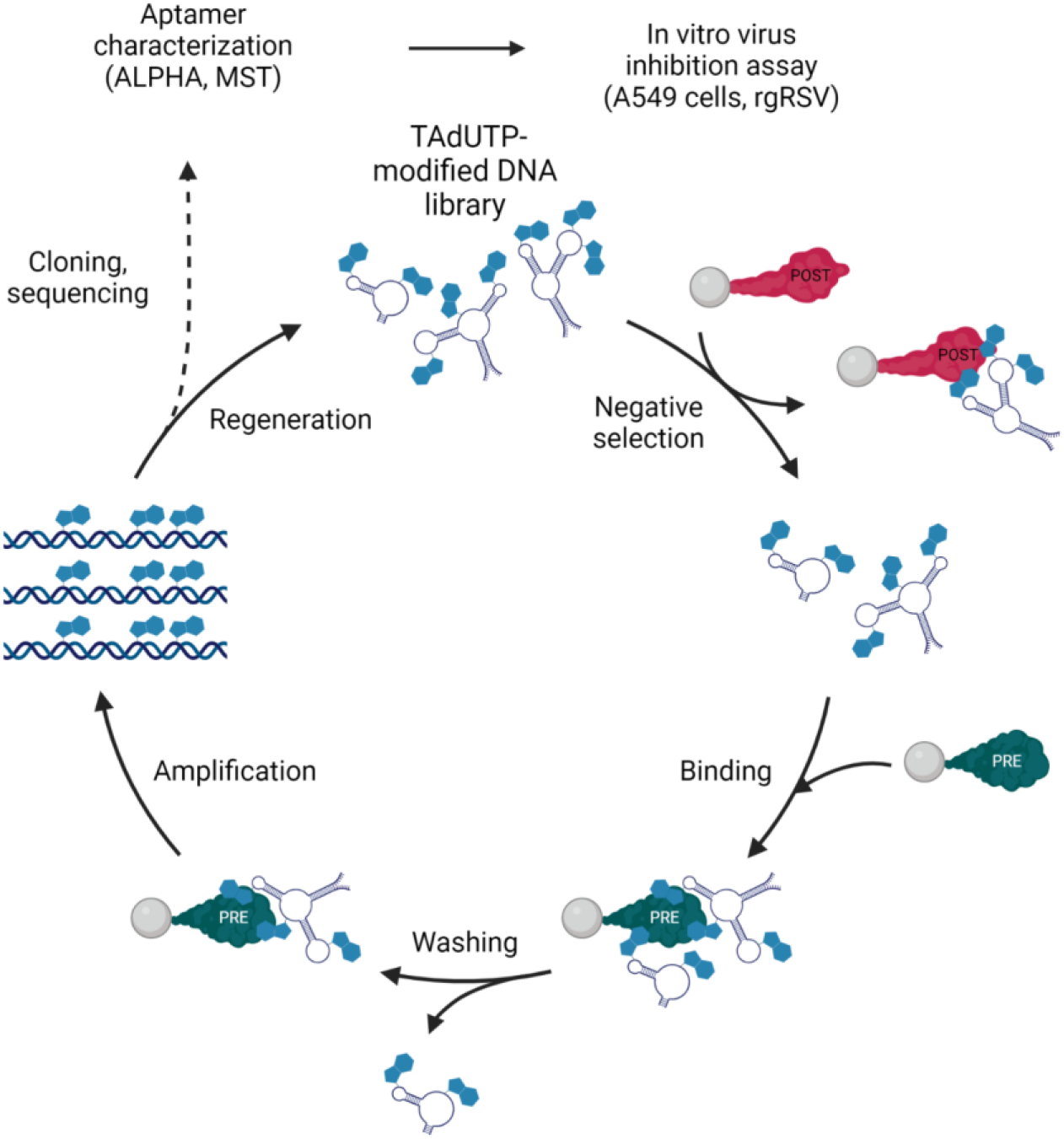
Schematics of RSV F protein selective aptamer generation and characterization. Created with BioRender.com.

### 2.2. Screening and characterization of aptamer candidates

Theoretically, target molecule binding oligonucleotides are enriched in successive cycles of SELEX. In practice, PCR bias also presents selection pressure on the amplified oligonucleotides; thus, the isolated aptamer candidates do not necessarily possess high affinity to the target molecule (17). The MEME analysis did not identify any common motif of the selected oligonucleotide that could have hinted the most auspicious aptamer candidates. Therefore, we set out to screen all sequenced oligonucleotides by Amplified Luminescent Proximity Homogeneous Assay (ALPHA).

To this end, we synthesized biotin labelled aptamer candidates by primer-blocked asymmetric PCR (PBA-PCR (33)) and mixed them with the prefusion form of F protein and palivizumab, and antibody which binds both prefusion and postfusion form of F protein. The biotinylated aptamers and the antibody were immobilized on Streptavidin-coated donor and Protein A-covered acceptor AlphaScreen beads, respectively. Out of the 70 aptamers, almost all showed binding to the target molecule and 14 aptamers produced 10-50 times higher fluorescence signal upon binding the prefusion form of F protein in comparison to both the F protein and the antibody minus controls (Figure S2). The measured elevated fluorescence intensities indicated success of aptamer selection. Of note, although several aptamer candidates possessed almost identical nucleotide compositions (see Supplemental data file), they demonstrated different affinities for the F protein. These data and the lack of common motif of oligonucleotides suggest that unique nucleic acid sequences are inevitable for the F protein-aptamer interaction.

Next, the protein F conformation discriminating capacity of the outstanding 14 aptamers was studied. We blended the aptamers either with the prefusion or the postfusion F protein at different concentrations applying the above-described experimental setting. All analysed aptamers produced an approx. 2 times higher relative fluorescence signal when mixed with the prefusion form (Figure 2) than upon incubating with the postfusion form of protein F.

**Figure 2.**
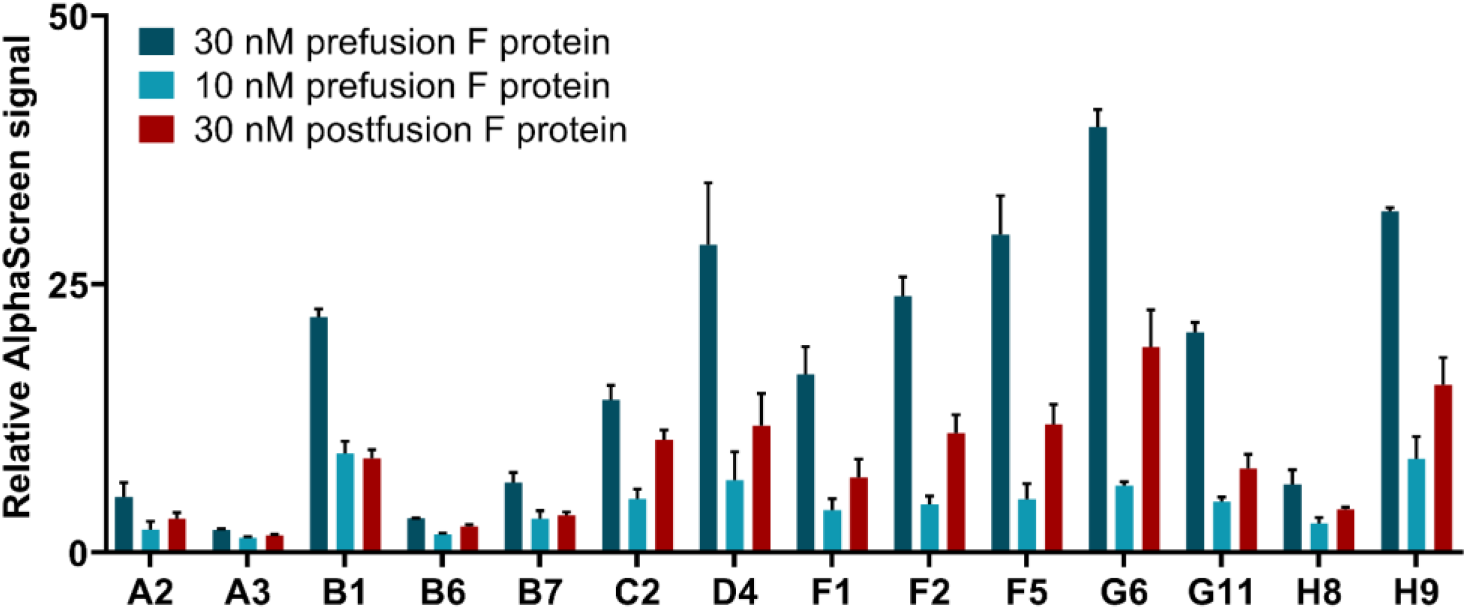
Determining the ability of modified aptamers to distinguish between RSV’s prefusion and postfusion F protein by AlphaScreen. Biotin labelled aptamers were mixed with either the prefusion or postfusion form of F protein and Palivizumab. Relative AlphaScreen signal was calculated by forming a ratio of the sample fluorescence and the aptamer-free background fluorescence. The increased values indicate selective binding of aptamers to the different F protein forms.

To analyse binding kinetics of aptamer-protein complex formation, we applied Microscale Thermophoresis (MST), a generally leveraged method for the analysis of interactions between biomolecules in solution. In MST experiments, the concentration of the smaller, fluorescently labelled molecule was kept constant, while a serial dilution of the larger, unlabelled interaction partner was added to the mixture.

The obtained data showed variability in the K_D_ values of different aptamer-protein F complexes. Some of the aptamers bound to their target with low μM dissociation constant (B1, C2, F5, G6, and H8), while the majority of them presented sub μM values, and the most eminent candidates formed aptamer-prefusion protein F complexes which possessed few 100 nM dissociation constants (A2, D4, H9). Of note, all studied aptamers demonstrated a much stronger affinity to the prefusion form than to the postfusion form of the target protein implying the efficiency of counter-selection steps of SELEX (Table S1).

It could seem contradictory that results of AlphaScreen and MST measurements were not in full harmony, i.e., the highest AlphaScreen signal producing aptamers did not consistently possess the lowest K_D_ values. However, this phenomenon was not entirely unexpected and could be explained by the inherent differences of the two interaction characterising methods. AlphaScreen requires the immobilization of the interaction partners, meanwhile MST determines the interaction between two partners in free solution.

In order to demonstrate the specificity of the most promising aptamer candidates (A2, B1, D4, G6, H8, H9), we embarked on measuring their putative interaction with various human pathogen virus proteins, i.e., human metapneumovirus fusion protein (MPV, variant v3B) (34), human parainfluenza virus type 3 fusion protein (PIV3, Q162C-L168C, I213C-G230C, A463V, I474Y variant) (35), SARS-CoV-2 spike protein (S-2P variant) (36), influenza H1 hemagglutinin protein (S106 DS26r) (37) using MST. The collected data (Figure S3, Table S2) demonstrated the selectivity of our aptamers.

### 2.3. Modified aptamers possess antiviral effect on A549 cells similar to that of palivizumab

Following in vitro characterization of the selected aptamers, we embarked on studying their virus infection inhibiting potential. Recombinant green fluorescent protein (GFP)-expressing RSV (rgRSV) carries the GFP gene prior to the full-length genome of RSV thus the infection is conveniently traceable by measuring fluorescence of the treated cells (38). First, to determine the optimal experimental conditions, a monolayer of A549 cells were infected with recombinant RSV expressing green fluorescent protein (rgRSV) and the fluorescence intensity were measured for 72 hrs. The obtained data showed that the signal peaked at 48 hrs; thus, this incubation time was applied in the ensuing experiments (Figure S4). In these treatment conditions, the modified aptamers remained relatively intact as opposed to the non-modified aptamers (Figure 3).

**Figure 3.**
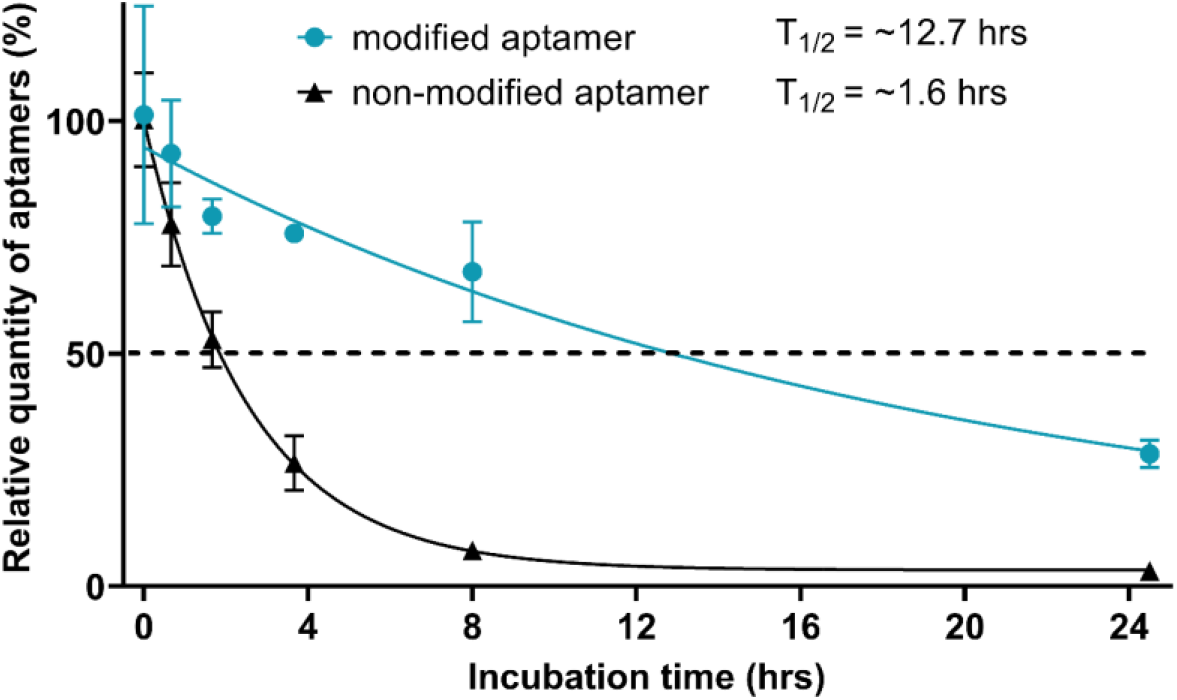
Stability of a modified and a non-modified aptamer was demonstrated by incubating A549 cells with aptamer infused growth media. The oligonucleotide concentration was determined by real-time PCR at the indicated time points. The degradation of aptamers was remarkably different, the modified aptamer seems to possess an approx. 8 times longer half-life in comparison to its non-modified variant.

Next, we chose 6 aptamers for viral inhibition studies, two of them, D4 and H9, were amongst the best performers both in AlphaScreen and MST measurements; A2 and G6 were outstanding according to AlphaScreen and MST studies, respectively; B1 and H8 were selected randomly from the previously analysed 14 aptamers (Figure S6). RgRSV was preincubated with either palivizumab or aptamers and added to A549 cells. Following the removal of rgRSV and washing the cells, the virus infected cell generated fluorescence signal was determined for 48 hrs. Both palivizumab and modified aptamer preincubated samples showed a strong decrease in fluorescence signal in comparison to control samples without palivizumab and aptamer. Importantly, the non-relevant or non-modified nucleotide holding aptamer (Table S3) preincubated rgRSV mixtures produced fluorescence intensity that were similar to controls without antibody or aptamer (Figure 4A). To better assess the antiviral effect of the aptamers, the area under the curve (AUC) was calculated from obtained growth curves, and it suggested at least a 50% decrease of infection upon treatment of cells with either palivizumab or aptamer preincubated rgRSV (Figure 4B). To provide further evidence for the reduction of virus infection, we applied a very different methodological approach, that is, the measurement of the amount of viral RNA by rt-qPCR. The rt-qPCR provided data were in concert with the AUC calculations thus corroborated the inhibitory effect of aptamers on RSV infection (Figure 4C).

**Figure 4.**
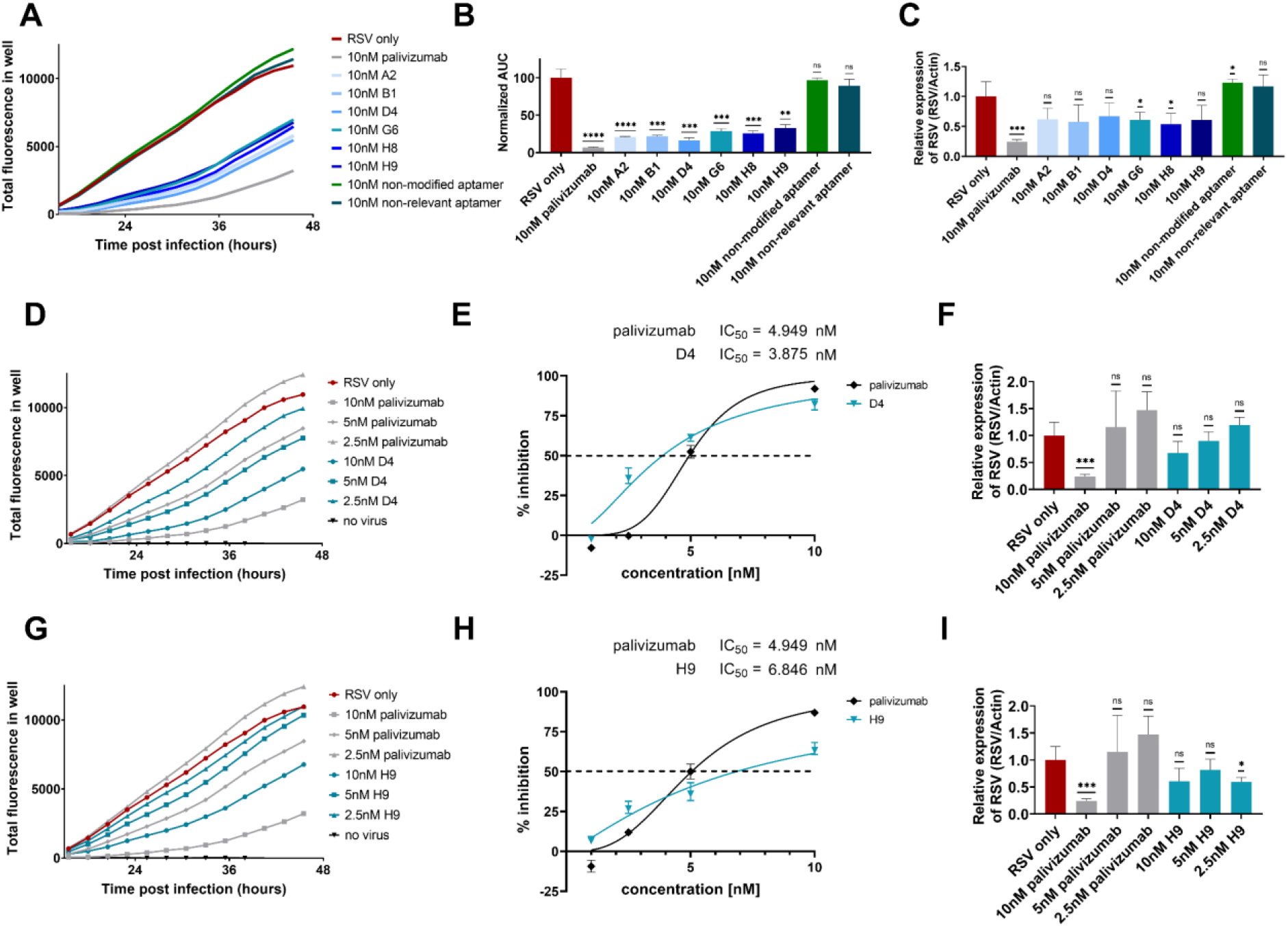
Modified aptamers exhibit antiviral effect upon RSV infection. Total fluorescence (TF) was measured of the rgRSV infected A549 cell culture (MOI of 1). Palivizumab or the aptamers (A2, B1, D4, G6, H8, H9, non-modified and non-relevant aptamers) were pre-incubated with rgRSV prior to infection. (A) Lower TF is measured when rgRSV is pre-treated with modified aptamers or palivizumab in comparison to infection with mock-treated RSV. The non-modified aptamer or a non-relevant aptamer had marginal effect on virus neutralization. (B) Area under the curve (AUC) was calculated, mean and standard deviations of three replicates are shown. (C) Rt-qPCR of the viral RNA in infected A549 cells verifies the antiviral effect of modified aptamers (D, G). The most promising aptamer candidates, D4 and H9, demonstrate the highest virus neutralizing capability. (E, H) Percent inhibition (calculated from the AUC) compared to the mock-treated rgRSV control. RgRSV infection is reduced by palivizumab and D4 or H9 in a very similar, concentration dependent manner. Error bars indicate SD, the dashed lines indicate 50% inhibition. (F, I) Reduction in the amount of viral genome detected in the infected A549 cells also signifies the antiviral effect of the modified aptamers. P-values were calculated using unpaired t-test (*P < 0.05, **P < 0.005, ***P < 0.001, ****P < 0.0001, ns = not significant).

After demonstrating that our SELEX protocol resulted in modified aptamers that could effectively restrict RSV infection at nM concentrations, we embarked on further studies of the outstanding aptamers. A dose response assay was carried out with a series of concentrations of aptamers that performed equally well both in ALPHA and MST (D4 and H9). The dose-dependent decrease in the growth curves of rgRSV in samples which were preincubated with the modified aptamers showed that both D4 and H9 efficiently protect against rgRSV infection. The half maximal inhibitory concentration (IC_50_) values of both aptamers were comparable to those of palivizumab; they fell in the low nanomolar range (Figure 4 D, E, F and G, H, I). Dose response of the other four aptamers and non-modified versions of D4 and H9 were also studied (Figure S7). Interestingly, G6 aptamer, which produced highest signal in AlphaScreen but possessed low affinity according to the MST measurement, also exhibited similar virus neutralizing property to that of Palivizumab.

Lastly, considering the most striking cytopathologic effect of RSV infection, we examined syncytia formation of A549 cells. The light and fluorescence microscopy observations also confirmed the above findings since the syncytia formation was equally reduced in wells where rgRSV was preincubated either with palivizumab or with D4 and H9 aptamers (Figure S8). Altogether, the RSV-inhibitory effect of selected aptamers has been demonstrated with three distinctly different approaches.

### 2.4. Modified aptamers have no effect on the viability of A549 cells

Considering the therapeutic potential of the aptamers, we set out to test the putative cytotoxicity of modified aptamers. A549 cells were incubated with various modified, non-modified aptamers and palivizumab at 10 nM concentration for 48 hrs and their viability was assessed by a resazurin-based approach. The fluorescence measurements showed that neither the palivizumab nor any of the aptamers had detectable influence on cell viability (Figure 5A). To determine the 50% cytotoxicity concentration (CC50), A549 cells were treated with a dilution range of one of the most promising aptamer candidates (D4) and a non-relevant aptamer. According to the obtained data, no significant viability change was detected even at the highest applied, 50 nM concentration (Figure 5B). These results indicate that cytotoxicity is not expected to hinder therapeutic application of modified aptamers.

**Figure 5.**
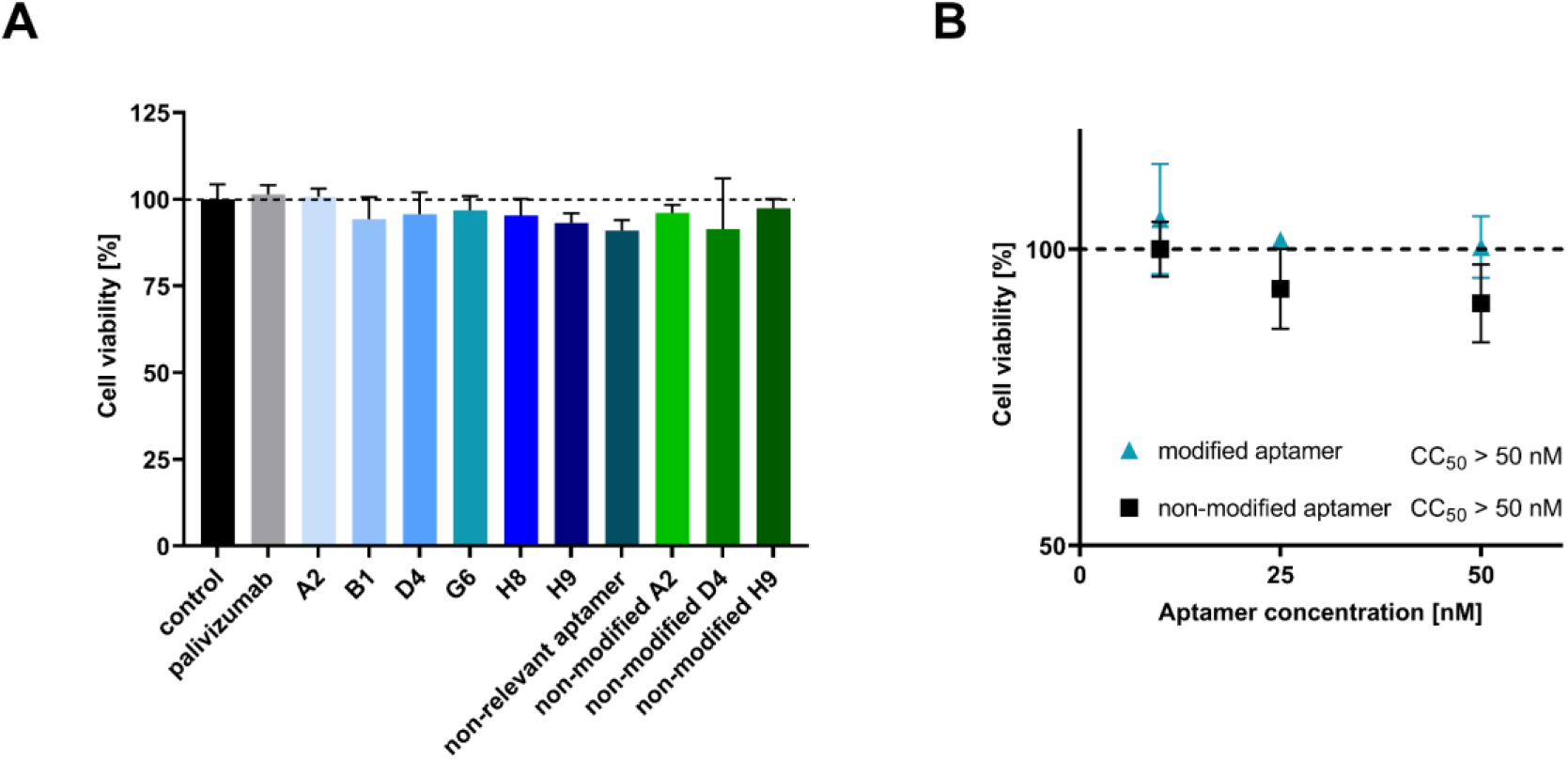
(A) Viability of A549 cells treated with aptamers and palivizumab. Cells were incubated with 10 nM of aptamers or palivizumab for 48 hrs, then their viability was assessed. Neither Palivizumab, nor any of the aptamers had a significant detrimental effect on the cell viability after 48 hrs. (B) Dose-dependent effect of one of the most promising modified aptamer candidates and a non-relevant aptamer on the viability of A549 cells. Cells were incubated with 50, 25 and 10 nM of modified aptamer or non-relevant aptamer for 48 hrs, then their viability was assessed. Neither the modified aptamer, nor the non-relevant aptamer had a detrimental effect on the cell viability after 48 hrs.

### 2.5. TAdUTP augments nuclease resistance of aptamers

Application of aptamers is frequently hindered by omnipresent nucleases. Therefore, we assessed durability of our modified nucleotide holding aptamer by challenging it with a lung epithelial cell line. A549 cells were incubated either with D4 aptamer or with its variant in which TAdUTP was replaced by dUTP, and the concentration of the oligonucleotides were determined by real-time PCR (Figure 3). Calculation of the relative quantities of the oligonucleotides clearly demonstrated elevated stability of the modified D4 aptamer. Approximately a third of the modified aptamer was present even after 24 hours incubation, while only a few percent of the original concentration was detectable in the case of TAdUTP lacking the oligonucleotide. The calculated half-life of the two oligonucleotides also confirmed the observed stability differences, i.e., reaching half of the original modified aptamer concentration took 12.7 hrs that is roughly eight times longer than the those seen with the non-modified nucleotide possessing oligonucleotides. These data suggest that aptamers with TAdUTP could withstand prevailing conditions of the airways for many hours.

## 3. DISCUSSION

Although a battery of virus selective aptamers has been generated, only a minority of them hold modified nucleotides and only two of them have been tested as potential drug candidates. Both are RNA aptamers and 2’-hydroxyl, -fluoro and -methyl modifications were applied to increase their nuclease resistance. A further common characteristic of these aptamers is that both target specific internal (unexposed) enzymes of Hepatitis C Virus and Japanese Encephalitis Virus, i.e., the NS5B replicase and methyltransferase, respectively (19,39). The consequence of targeting proteins that are not surface exposed and only expressed intracellularly is the requirement of internalization of the aptamers into the cells for blocking virus replication. Indeed, these aptamers were driven into the cells either by lipofectamine-based transfection or by cholesterol conjugation.

We consider the therapeutic application of virus surface protein binding aptamers more straightforward and potentially more effective; therefore, a stabilised variant of prefusion F protein was utilized as a target of SELEX. To increase the success rate of selection and durability of aptamers in physiological conditions, a base modified DNA oligonucleotide library with tryptophan-like side chains was used during each step of the SELEX procedure. Furthermore, considering that the most effective RSV neutralizing antibodies are known to bind prefusion F specific antigenic sites, the postfusion form of protein F was used either as a competitor within the binding phase of selection or as an immobilised target of counter-selection steps. Application of the modified nucleotide and the stringent counter selection resulted in aptamers that favour the prefusion over the postfusion form of F protein thus are expected to have enhanced virus infection inhibiting capability. Interestingly, the modified nucleotide was enriched in all studied aptamers hinting the importance of the indolyl group rendered hydrophobicity in the aptamer-protein complex formation. Presumably, the modified nucleotides enabled the interaction between the aptamer and the hydrophobic amino acid stretch of the prefusion F specific antigenic site Ø.

Sanger sequencing and *in silico* analysis of the SELEX obtained oligonucleotides resulted in identification of a panel pf singletons and nucleic acid sequences which were present in three copies but no common motif of the oligonucleotides could be identified. Since the computational analysis did not provide obvious data to guide the identification of the most promising aptamer candidates, all identified oligonucleotides were examined by *in vitro* methods. Two generally leveraged homogenous assays, ALPHA and MST, were applied for screening and affinity determination of oligonucleotides, respectively. The two approaches provided partially contradictory results, i.e., the best performers of ALPHA measurements were not identical with the lowest K_D_ possessing aptamers. This phenomenon is however not unprecedented; it has been described that the various methods could provide dissimilar result for the very same molecular interactions (40). These results highlight the importance of applying different methods for identification of the prime aptamer candidates.

*In vitro* screening and characterisation of the selected oligonucleotides is a crucial step of aptamer generation but the data provided by these approaches have to be handled cautiously. Analysis of the virus inhibitory potential of the six aptamers demonstrated that the virus infection blocking capacity of the aptamers is not necessarily proportional to the AlphaScreen or MST data. According to the virus infection generated fluorescence and the viral genome quantifying RT-qPCR measurements, all studied aptamers prevent the virus infection with a comparable efficiency. This signifies that even though *in vitro* interaction assays narrow down the number of potential aptamer candidates for further analysis, the practical applicability of the aptamers has to be tested in the conditions of their intended application.

The implemented dose response assays demonstrated that the low nanomolar half maximal inhibitory concentration values of our aptamers are comparable to those of palivizumab. These data encouraged us to study those attributes of the aptamers which could hinder their therapeutic application. It has been described that phosphorothioate and 2’ fluoro modified nucleotide composed oligonucleotides could lead to cell death by triggering the p53 pathway (41,42). Our analysis showed no cytotoxicity of the aptamers even following 48 hrs incubation at ten times of IC_50_ oligonucleotide concentration. Although we did not study activation of the apoptotic pathways, these results indicate that TAdUTP modified oligonucleotides do not have detrimental effects on the cells.

A further bottleneck of clinical application of aptamers is their nuclease sensitivity. It has previously been reported that the incorporation of hydrophobic modifications at 5-position of pyrimidine nucleotides imparts a substantial increase in resistance to degradation in human plasma (43). Our results are in accordance with these findings; introduction of the indolyl group to the 5-position of dUTP resulted in about eight times longer half-life of the oligonucleotide in lung epithelial cell culture. Of note, our studied oligonucleotides were not protected against exonucleases; therefore, a simple 3’capping such as addition of an inverted nucleotide is expected to enhance further stability of TAdUTP modified aptamers. We believe that the presented results testify that virus surface protein targeting modified aptamers could yield efficient and cost-effective drug candidates, which could keep pace with the continuously evolving pathogenic viruses.

## 4. MATERIALS AND METHODS

### 4.1. Generation of tryptamino-modified aptamer library

The modified nucleotide, 5-[(3-Indolyl)propionamide-N-allyl]-2’-deoxyuridine-5’-triphosphate (TAdUTP) and CleanAmp dATP, dCTP and dGTP was purchased from TriLink. To generate the tryptamino-modified library, a water-in-oil emulsion PCR reaction were set up by using 1x KOD XL reaction buffer, 0.2mM dNTP mixture (containing dATP, dCTP, dCTP, TAdUTP in 2.5 mM concentration each), 2.5 U of KOD XL, 40 μM and 200 nM final primer and template concentrations, respectively. The PCR mixture was emulsified according to the manufacturer’s protocol (Micellula DNA Emulsion & Purification Kit (Roboklon)). PCR products from the emulsion were recovered using OligoClean & Concentrator Kit (ZymoResearch) according to the manufacturer’s protocol. The applied primers and oligonucleotides were synthesised by IBA, the detailed sequences can be found in Table S1. Amplification conditions were: 3 min denaturation at 95 °C, 7 cycles of 95 °C for 30 s, 60 °C for 5 s, 72 °C for 30 s, and a final extension at 72 °C for 3 min. The PCR products were analysed by 10 % polyacrylamide gel electrophoresis and 1 μL of GeneRuler Low Range DNA Ladder was used as molecular weight marker.

### 4.2. Production of prefusion and postfusion RSV F

Both prefusion (DS-Cav1) and postfusion forms of trimeric RSV F were produced by transient transfection, and purified by conventional chromatography as described previously.^10^ In both cases, final size-exclusion chromatography step was utilized to ensure trimeric homogeneity of the F protein.

### 4.3. Selection of RSV F protein selective modified aptamers

The SELEX conditions applied for the generation of RSV F protein selective aptamers is described in the Supplemental data file according to the MAPS guidelines (44). In short, the stabilised form of prefusion (“DS-Cav1”) and postfusion F protein was used as targets and negative targets of SELEX, respectively. 5.7 mg of beads from MACSflex™ MicroBead Starting Kit (Miltenyi Biotec) was reconstituted in 257 μl of Reconstitution Buffer. This bead was modified using 15 μg of prefusion or postfusion F protein in 25 μl of MES buffer by O/N incubation at 4°C. Success of immobilization was determined using SDS-PAGE and Coommassie-blue staining.

An aptamer selection was performed to generate RSV F protein selective aptamers using the generated tryptophan-like side chain holding aptamer library. The SELEX of RSV F specific modified aptamers were obtained by 7 iterative cycles with increasing selection pressure. The modified oligonucleotide library was heated to 95 °C for 5 min and immediately cooled on ice. First, approx. 1 nmole of the oligonucleotide library was incubated with non-modified Miltenyi beads in selection buffer (10 mg/L BSA, 0.02% Tween 20, 1 mg/L Salmon sperm DNA, 10 mg/L mucin in PBS) for 60 minutes using mild shaking to exclude the bead and mucin binding oligonucleotides. The F protein modified beads were incubated with the supernatant of the previous step for 30 minutes, then washed with 2 × 1000 μl PBS, followed by isolation of DNA pool using alkali elution and neutralization of protein F-bound oligonucleotides. Water-in-oil emulsion PCR was carried out to amplify the bound sequences using Micellula DNA Emulsion & Purification Kit (Roboklon). The PCR mixture contained KOD XL 1x reaction buffer, 2U of KOD XL polymerase (Toyobo) 0.4–0.4μM of untagged forward and biotinylated reverse primers of L8 library, and 0.1 mM each CleanAmp dATP, dGTP, dCTP mixed with TAdUTP. PCR products from the emulsion were recovered using OligoClean & Concentrator Kit (ZymoResearch) according to the manufacturer’s protocol. Amplification conditions were: 3 min at 94 °C, 25 cycles of 94 °C for 30 s, 60 °C for 5 s, 74 °C for 30 s and 74 °C for 5 min after the last cycle. The success of amplification was monitored by 10% polyacrylamide gel electrophoresis and staining with GelGreen DNA dye (Biotium).

For the regeneration of ssDNA by alkali denaturation, the PCR product was coupled to 25 μl streptavidin-coated paramagnetic beads (Dynabeads Streptavidin, M-280, Thermo Scientific) for 15 min and washed with 3 × 1000μl of PBS. The non-biotinylated strands were separated from the immobilized complementary strand by 10 min incubation with 50 μl of fresh 20 mM NaOH. The eluted ssDNA was immediately neutralized by addition of 7.5 μl of 200 mM NaH_2_PO_4_. In the following rounds of selection, the postfusion form of F protein was used either coupled to paramagnetic beads as a counter-target molecule (in rounds no. 4, 6, and 7) or the selection buffer was complemented with an excess of postfusion F (5-10 times in excess to the prefusion form in round 2, 3, 5, and 7). To further increase the selection pressure, the incubation conditions and washing steps were changed. In round 4, 5, 6, 7 the incubation time was reduced to 25, 20, and 15 min, respectively. In the third round an additional washing step with 1 ml of 0.15 mM dextran-sulphate in PBS (pH 7.4) was also introduced, in the fourth round, the bound sequences were challenged by washing the beads with 1 ml of 18 μM L-tryptophan solution as well. In the final selection round, the incubation time was 15 min and the beads were washed twice with 1 ml of 0.3 mM dextran-sulphate solution, twice with 1 ml of 18 μM L-tryptophan solution and twice with PBS solution. The PCR product of the last selection step was inserted into a cloning vector (Zero Blunt TOPO PCR Cloning Kit, Thermo Fischer Scientific) and transformed into One Shot™ TOP10 Chemically Competent E. coli cells (Invitrogen). 130 of the colonies were analysed by colony PCR (using an M13 primer set, Table S1) and capillary electrophoresis (LabChip GX, PerkinElmer) using the DNA 1K Reagent Kit with DNA HT 5K LabChip single sipper chip. For the latter, the colony PCR products were diluted 40x in TE buffer. The sequences of colony PCR products of correct size were determined by Sanger sequencing.

### 4.4. Generation of modified aptamers

AlphaScreen requires biotinylated aptamers which were generated by PBA-PCR (primer blocked asymmetric PCR). The 25 μl PCR mixture contained KOD XL 1x reaction buffer, 2 U of KOD XL polymerase, 0.5 μM of biotinylated forward primer, 25 nM untagged reverse primer, 475 nM 3’-P reverse primer, 0.2 mM each CleanAmp dATP, dGTP, dCTP mixed with TAdUTP, 0.5 μl of 40x diluted colony PCR template. Amplification conditions were: 3 min at 94 °C, 45 cycles of 94 °C for 30 s, 60 °C for 5 s, 74 °C for 30 s and 74 °C for 5 min after the last cycle. Following PCR, the complement of the reverse primer was added at 5 μM concentration, the mixture was heated to 95 °C for 10 minutes.

Cy5-labelled modified and non-modified aptamers were generated for MST and virus neutralization assays. For the modified aptamers, the 50 μl PCR mixture contained KOD XL 1x reaction buffer, 2 U of KOD XL polymerase, 1 μM of Cy5-labelled forward primer and biotin-labelled reverse primer, 0.2 mM each CleanAmp dATP, dGTP, dCTP mixed with TAdUTP, 0.5 μl of 40x diluted colony PCR template. For the non-modified aptamers, a dUTP mix (Bioline), containing dATP, dGTP, dCTP and dUTP, was used as substrate for the reaction. Amplification conditions were: 3 min at 94 °C, 30 cycles of 94 °C for 30 s, 60 °C for 5 s, 74 °C for 30 s and 74 °C for 5 min after the last cycle. Single-stranded DNA was regenerated using akali denaturation as described in “Selection of RSV F protein selective modified aptamers”.

The success of amplification was monitored by 10% polyacrylamide gel electrophoresis and staining with GelGreen DNA dye.

### 4.5. Screening of aptamers

The interaction assays were performed using white 384-well Optiplates (PerkinElmer) in total volume of 18 μl using Protein A acceptor and Streptavidin donor beads (PerkinElmer). Varying amounts of the prefusion and postfusion form of RSV F protein in PBS supplemented with mucin (10 mg/L), BSA (1 mg/ml), and Salmon sperm DNA (1 mg/L) were incubated with 10 nM final concentration of modified biotinylated aptamer and protein F selective antibody. Following 20 min incubation at RT, the acceptor and donor beads were added at 20 mg/L final concentrations in two steps. First, the Protein A acceptor beads were added and incubated for 30 minutes at RT and that was followed by the addition of Streptavidin donor beads and further 30 min incubation. Fluorescent signal was detected by using an EnSpire multilabel plate reader (PerkinElmer).

### 4.6. Microscale thermophoresis

10, 15 or 20 nM of Cy5-labelled aptamers were mixed with a 16-fold, 1:1 serial dilution of protein (human metapneumovirus fusion protein (MPV, variant v3B), human parainfluenza virus type 3 fusion protein (PIV3, Q162C-L168C, I213C-G230C, A463V, I474Y variant), SARS-CoV-2 spike protein (S-2P variant), influenza H1 hemagglutinin protein (S106 DS26r). All mixtures were prepared in 0.05% Tween 20 in PBS and the measurement was carried out in Monolith NT™ Standard capillaries (NanoTemper Technologies). Excitation power and MST power of Microscale Thermophoresis instrument (Monolith NT.115, NanoTemper Technologies) was set to 50% and 20%, respectively, the temperature was set to 25°C during the measurements. All obtained data was analysed using the MO.Affinity Analysis v2.3 software (NanoTemper Technologies), where dissociation constants were calculated from the fitted curve using Michaelis-Menten kinetics.

### 4.7. Cultivation of A549 cell line and rgRSV

A549 cells were cultured in complete media consisting of Dulbecco’s MEM (Gibco) modified with 10% FBS (Gibco) and 1% PenStrep (Gibco). Confluent cells were split every 4 days. RgRSV was cultured as described elsewhere.^28^

### 4.8. In vitro virus neutralization assay

A549 cells were seeded in a 96-well black clear bottom plate at 2.5 × 10^4^ cells/well in DMEM (10% FCS, 1% PenStrep (Gibco)) and cultured at 37°C, 5% CO_2_. At 24 hours post-seeding, cells were washed with sterile PBS two times and infected with rgRSV at multiplicity of infection (MOI) of 1. Dilutions were previously made from rgRSV in DMEM (1% PenStrep, omitting FCS), palivizumab and aptamers are added in a final concentration of 1, 2.5, 5 or 10 nM. Pre-incubation of antiviral agents and RSV were carried out for 1.5 hr at 37°C, 5% CO2. Prior to infection, A549 cells were washed 2 times with 100 uls of PBS, the pre-incubated samples were added and mixture were incubated for 1.5 hr at 37°C, 5% CO_2_. Infected cells were washed 2 times with PBS, then 100uls of DMEM (10%FCS, 1% PenStrep) was added to each well. Every treatment was carried out in triplicates. The plate was transferred to the humidity chamber of a fluorescent plate reader (Tecan Spark), and measurement of generated fluorescence was performed for 48 hrs at 37°C, 5% CO2. Infected cells were also imaged using fluorescence microscopy (Leica DMIL LED Inverted Routine Fluorescence Microscope). Cells were then lysed with iScriptTM RT-qPCR Sample Preparation Reagent according to the manufacturer’s protocol.

### 4.9. Rt-qPCR of RSV infected cells

Reverse transcription of the viral genome in the cell lysates was carried out using iScript™ Reverse Transcription Supermix for RT-qPCR. Then, qPCR of the cDNA was performed using a primer set for the N gene of RSV (45) and ACTB (Applied Biosystems, TaqMan Assay ID: Hs99999903_m1) as a housekeeping gene, mixed with and SsoAdvanced Universal SYBR Green Supermix or SsoAdvanced Universal Probe Supermix and measured by CFX real-time PCR system. QPCR cycling conditions were as follows: 95°C for 3mins, 50 cycles of 95°C for 10 seconds, 60°C for 20 seconds; followed by melting curve analysis. Data was analysed using CFX Maestro (BioRad).

### 4.10. Cell viability assay

A549 cells were seeded in a 96-well black clear bottom plate at 2.5 × 10^4^ cells/well in DMEM (10% FCS, 1% PenStrep (Gibco)) and cultured at 37°C, 5% CO_2_. At 24 hours post-seeding, cells were treated with a dilution series of modified aptamers. Cells treated with PBS were used as a negative control. After 48 h, the cell viability was assessed using the CellTiter Blue kit (Promega) according to the manufacturer’s instructions. Fluorescence was detected on the Spark Plate Reader (Tecan).

### 4.11. Assessment of the degradation of aptamers

A549 cells were seeded in a 96-well black clear bottom plate at 2.5 × 10^4^ cells/well in DMEM (10% FCS, 1% PenStrep (Gibco)) and cultured at 37°C, 5% CO_2_. At 24 hours post-seeding, cells were washed with sterile PBS two times. DMEM (10% FCS, 1% PenStrep) was mixed with the aptamers in the final concentration of 10 nM and added to A549 cells. Samples were taken from the supernatant of cells after 0 h, 0.5 h, 1 h, 2 h, 4 h, 8 h and 24 h of incubation, then snap frozen in liquid nitrogen immediately. Next, qPCR was carried out using QuantStudio 12 K Flex PCR System. The 10 μl PCR mixture consisted of 0.4 μl of 10 μM unlabeled reverse and forward primer each, 5 μl of qPCRBIO SyGreen Mix Lo-ROX (PCR Biosystems), 2 μl sample, 2.2 μl nuclease-free water. The reaction conditions of real-time qPCR were the following: initial denaturation for 2 min at 95 °C, followed by 40 cycles of denaturation for 5 s at 95 °C, annealing for 20 s at 60 °C. Melting curve analysis was performed from 60 °C to 95 °C.

### 4.12. Statistical analysis

All graphs were plotted, and statistical analyses were performed using GraphPad Prism (version 9.1.2.). Graphs represent the mean of three replicates ± standard deviation. P-values were calculated using unpaired t-test (*P < 0.05, **P < 0.005, ***P < 0.001, ****P < 0.0001, ns = not significant). Area under the curve (AUC) was calculated using the obtained fluorescence intensity 0 - 30 hours post-infection. The percentage of inhibition was calculated using the following formula: (1 - (AUC of sample / AUC of mock-treated rgRSV infected cells)) × 100. The inhibitory concentration of aptamers that reduced viral levels by 50% (IC_50_) was estimated by fitting a nonlinear curve of variable slope to the percentage of inhibition of samples.

## 5. DATA AVAILABILITY

The data underlying Figs. 2–5, as well as all Supplemental figures are enclosed in the Supplemental data file. Any additional data from this study is available from the authors upon reasonable request. Supplemental information is provided with this paper.

## 6. ACKNOWLEDGEMENTS

Funded in part by the Intramural Research Program of the Vaccine Research Center, NIAID, NIH (ZIA AI005024-22) and by the Hungarian National Research, Development and Innovation Office (NKFIH grant number: ANN-139564). Project no. TKP2021-EGA-24 has been implemented with the support provided by the Ministry of Innovation and Technology of Hungary from the National Research, Development and Innovation Fund, financed under the TKP2021-EGA funding scheme. K.P. was supported by a Federation of European Biochemical Societies (FEBS) Short-term fellowship.

## 7. AUTHOR CONTRIBUTIONS

T.M. and M.dJ. conceptualised this work; L.O. prepared Ds-Cav1 and postfusion F protein along with pre-fusion trimers from human metapneumovirus and human parainfluenza virus type 3; N.D. provided hemagglutinin trimer; A.S.O. provided SARS-CoV-2 spike; K.P. and T.M. designed and performed RSV F aptamer selection; K.P., Z.J.T. designed and performed aptamer characterization assays; K.P., M.E., M.dJ. designed and executed virus neutralization experiments; K. P. analysed and interpreted the results; T.M., K.P., M.dJ, P.D.K. wrote and edited the manuscript. All authors reviewed the results and approved the final version of the manuscript. The manuscript hasn’t been accepted or published elsewhere.

## 8. TRANSPARENCY DECLARATIONS

The authors declare no conflict of interest.

