## Supplemental information for "Tryptophan-like side chain holding aptamers inhibit respiratory syncytial virus infection of lung epithelial cells"

| Name | Abundance of sequence in selected aptamer pool | Nucleotide composition of the aptamer | | | | | Sequence (5'-3') |
|---|---|---|---|---|---|---|---|
| | | T% | G% | A% | C% | TAdU% | |
| A5 | 9,4% | 3,9% | 31,6% | 19,7% | 17,1% | 27,6% | GATCGAGCCAGCTACGTCAGGUAAGAUCAUGUGAUGUGUGUUGGUUUGUUGUGUGUGUGACAUGCCUCAGCGACUA |
| A7 | 4,2% | 3,9% | 38,2% | 17,1% | 15,8% | 25,0% | GATCGAGCCAGCTACGTCGAGUUUGAGUGUGUUAGGUGGGGUUGGUGGGUUGGGUUUAGACAUGCCUCAGCGACUA |
| A3 | 3,1% | 3,9% | 43,4% | 15,8% | 17,1% | 19,7% | GATCGAGCCAGCTACGTCGACAUGUAGGGUGGUGUGGUUGGGGGUGGUGUGGUGGGGUGACAUGCCUCAGCGACUA |
| B1 | 3,1% | 3,9% | 42,1% | 17,1% | 15,8% | 21,1% | GATCGAGCCAGCTACGTCAGUUAUGGUAUAGGUUGGUGGUGGGGGGGGGUUGUGGUGGGACAUGCCUCAGCGACUA |
| B2 | 3,1% | 3,9% | 30,3% | 17,1% | 17,1% | 31,6% | GATCGAGCCAGCTACGTCAGUUGUGUAUGUGGUGUAUGUUGGGUUUGUUGUUCGUUUAGACAUGCCUCAGCGACUA |
| B6 | 3,1% | 3,9% | 42,1% | 13,2% | 17,1% | 23,7% | GATCGAGCCAGCTACGTCUGUAUGGGUGUGGUGUGUGGGGGUGGGUGGUGGGUUGUUCGACAUGCCUCAGCGACUA |
| B7 | 3,1% | 3,9% | 42,1% | 18,4% | 15,8% | 19,7% | GATCGAGCCAGCTACGTCAAUAGGGUGUGAUGUGGUUGGGGGGUAGGUGGGUGGGUGUGACAUGCCUCAGCGACUA |
| A4 | 2,1% | 3,9% | 27,6% | 17,1% | 21,1% | 30,3% | GATCGAGCCAGCTACGTCUAAUGUAUGUGUGUGUUUGUGUGUUUCCUACUGUGUGUGCGACAUGCCUCAGCGACUA |
| D9 | 2,1% | 3,9% | 42,1% | 15,8% | 17,1% | 21,1% | GATCGAGCCAGCTACGTCGACAUGUAGGGUGGUGUGGUUGGGGGUUGUGUGGUGGGGUGACAUGCCUCAGCGACUA |
| E5 | 2,1% | 3,9% | 36,8% | 15,8% | 18,4% | 25,0% | GATCGAGCCAGCTACGTCAUGUCUGGGUGGGUGAGGGUGGGUUAUGUGUGCUUUGUGUGACAUGCCUCAGCGACUA |

**Supplementary figure** 1 Overrepresentation of the modified nucleotide and guanosine in the most abundant sequences in the selected aptamer pools.

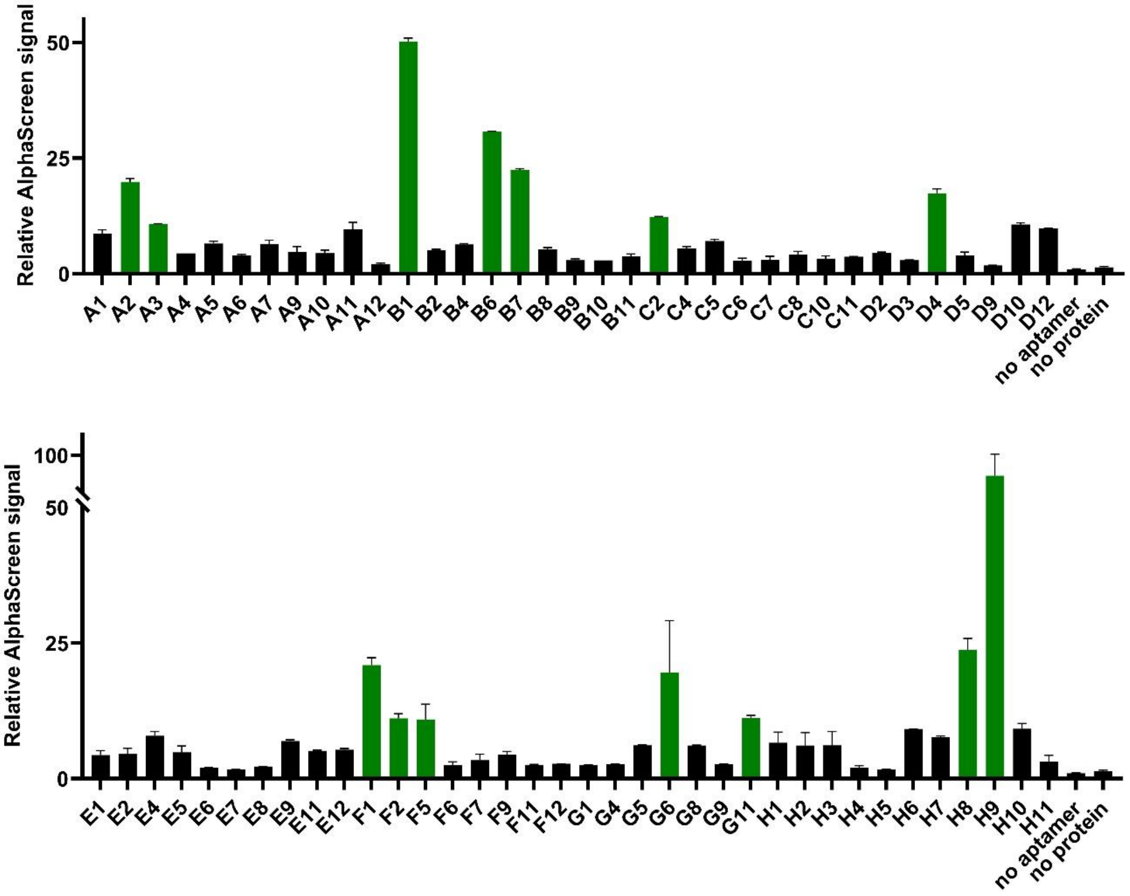


**Supplementary figure** 2 Screening of modified aptamers using AlphaScreen. Biotin-labelled aptamers were mixed with 30 nM of prefusion form of F protein and Palivizumab. An increased fluorescence signal is detected upon binding of aptamers to prefusion F protein. Aptamers A2, A3, B1, B6, B7, C2, D4, F1, F2, F5, G6, G11, H8 and H9 (indicated as green bars) were chosen for further analysis. Relative AlphaScreen signal was calculated by forming a ratio of the sample fluorescence and the aptamer-free background fluorescence. The increased values indicate selective binding of aptamers to prefusion F protein.

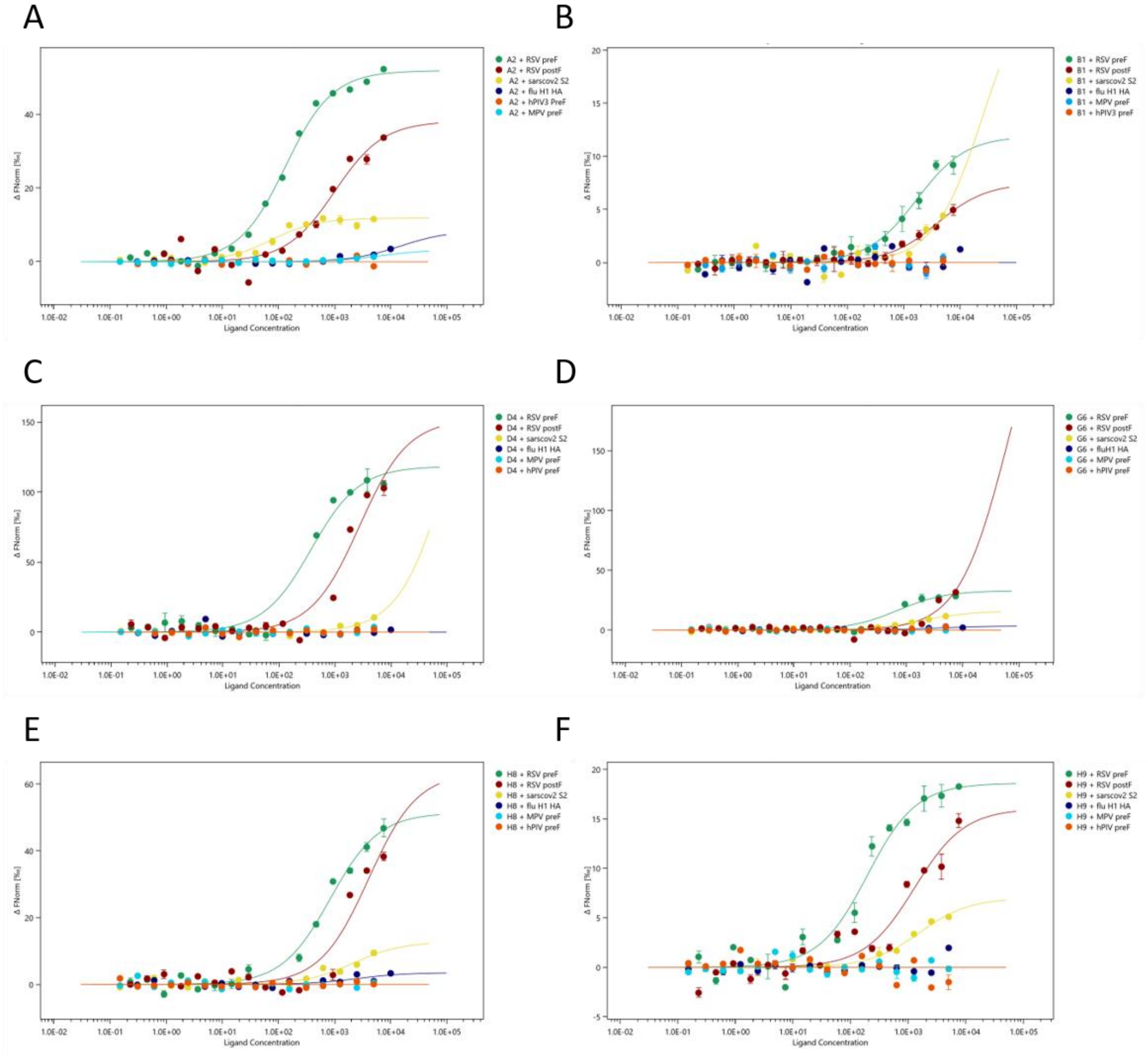


**Supplementary figure 3** Selectivity of the 6 functionally analysed aptamers was determined by MST. A serial dilution was made of viral proteins (RSV F prefusion protein (Ds-Cav1), RSV F postfusion, SARS-Cov2 spike protein (Sarscov2 S2), influenza H1 hemagglutinin (fluH1 HA), human parainfluenza virus prefusion protein (hPIV preF) and human metapneumovirus prefusion protein (MPV preF) and mixed with a constant concentration of Cy5-labelled modified aptamers, then the IR induced thermophoresis was measured. The calculated $K_D$ values are listed in Supplementary table 2.

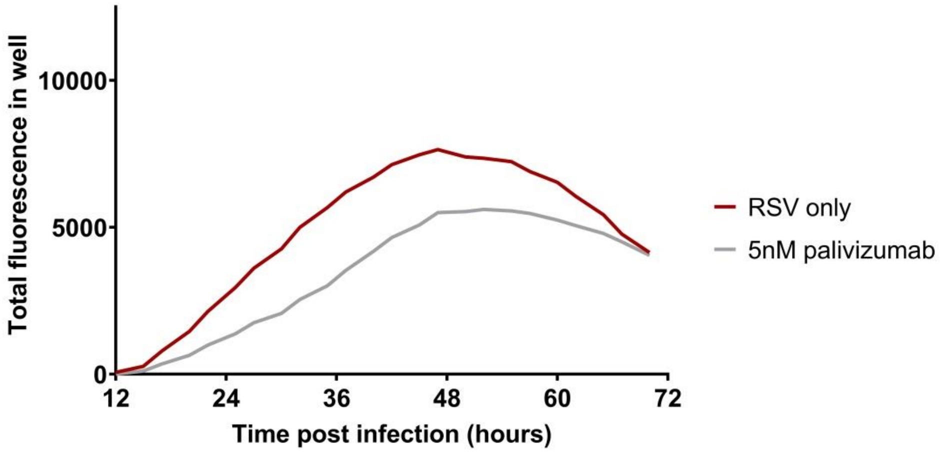


**Supplementary figure 4** A549 cells were infected with rgRSV and the infection rate, proportional to the fluorescence signal, was measured up 72 hrs.

| Name | Abundance of sequence in selected aptamer pool | Nucleotide composition of the aptamer | | | | | Sequence (5'-3') |
|---|---|---|---|---|---|---|---|
| | | T% | G% | A% | C% | TAdU% | |
| **A2** | 1,04% | 3,9% | 35,5% | 18,4% | 18,4% | 23,7% | GATCGAGCCAGCTACGTCAUGCUGUCAAUGGGUGGGUGUGGUGGGGUUGAUGUUAUGUGACAUGCCUCAGCGACUA |
| **B1** | 3,13% | 3,9% | 42,1% | 17,1% | 15,8% | 21,1% | GATCGAGCCAGCTACGTCAGUUAUGGUAUAGGUUGGUGGUGGGGGGGGGUUGUGGUGGGACAUGCCUCAGCGACUA |
| **D4** | 1,04% | 4,0% | 41,3% | 18,7% | 16,0% | 20,0% | GATCGAGCCAGCTACGTCAUGGUUGGGAGGUGUUGUGGGGAUGGUGGGUGGGAGUAU-GACAUGCCUCAGCGACUA |
| **G6** | 1,04% | 3,9% | 36,8% | 23,7% | 15,8% | 19,7% | GATCGAGCCAGCTACGTCAUGGGUAAAGGGAGGGUGUAGUGUAUAUGUGUGGUGGGAUGACAUGCCUCAGCGACUA |
| **H8** | 1,04% | 3,9% | 42,1% | 15,8% | 15,8% | 22,4% | GATCGAGCCAGCTACGTCAUGUGGGGUGUGGUGGUGGGGGGUUGUGUGGUGGGUUAAUGACAUGCCUCAGCGACUA |
| **H9** | 1,04% | 3,9% | 42,9% | 15,6% | 15,6% | 22,1% | GATCGAGCCAGCTACGTCAUGUUAAGGUGGUGGUUUGUGGGGGGGGUGUGGUGGGUGUGACAUGCCUCAGCGACUA |

**Supplementary figure** 5 Sequence, abundance and nucleotide composition of the 6 aptamers chosen for viral inhibition studies.

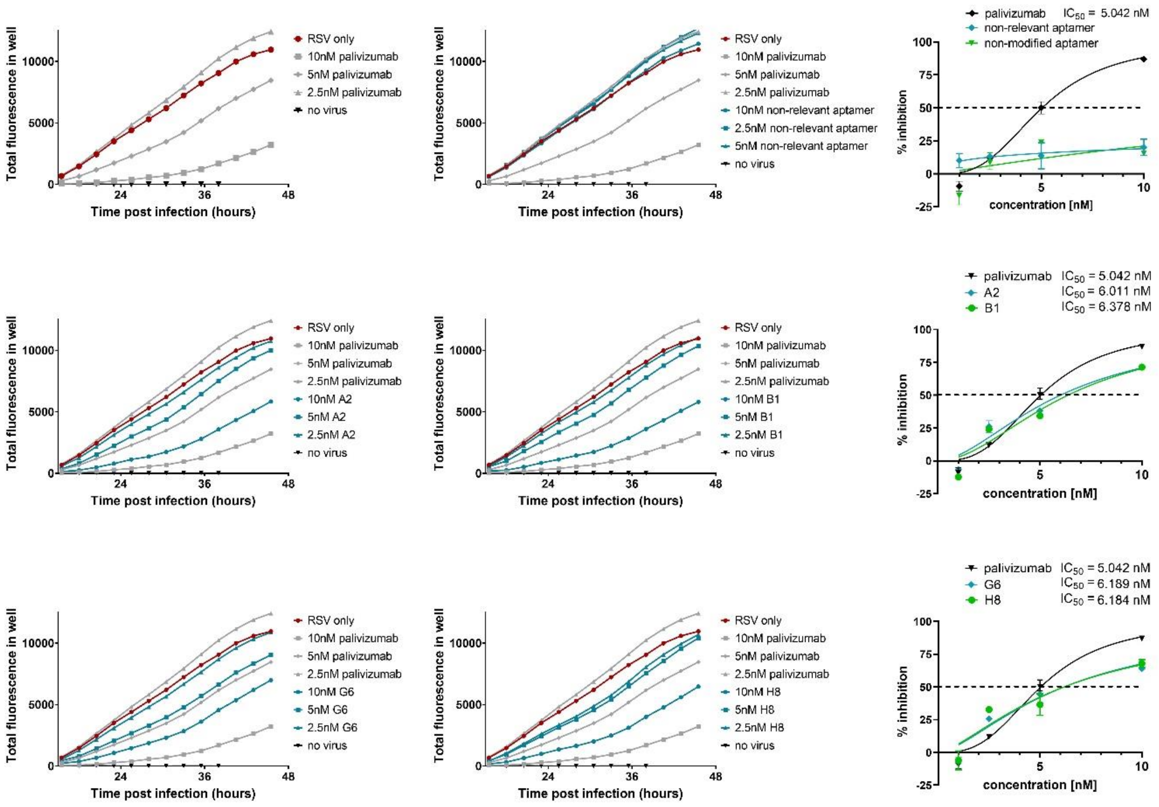


**Supplementary figure 6** Total fluorescence (TF) measurement of the rgRSV infected A549 cell culture. Different concentrations of palivizumab and the aptamers (A2, B1, G6, H8, the non-modified and the non-relevant aptamer) were pre-incubated with rgRSV prior to infection. Lower TF is measured when rgRSV is pre-treated with modified aptamers or palivizumab in comparison to infection with mock-treated RSV. The non-modified version of H9 and the non-relevant aptamer had marginal effect on virus neutralization (calculated from the AUC) compared to the mock-treated rgRSV control. RgRSV infection is reduced by palivizumab and by all four modified aptamers in a very similar, concentration dependent, manner. Error bars indicate SD, the dashed lines indicate 50% inhibition.

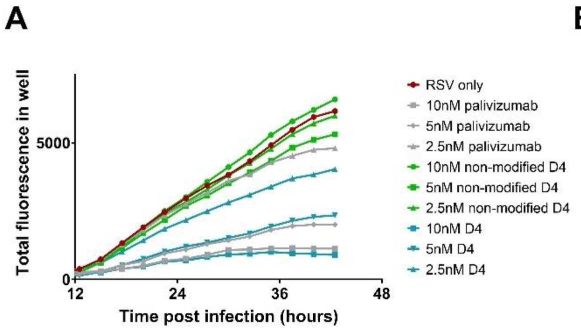


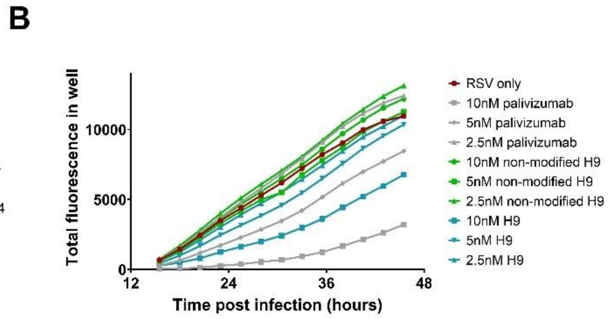


**Supplementary figure 7** Total fluorescence (TF) measurement of the rgRSV infected A549 cell culture. Palivizumab, the most promising aptamers (D4, H9) or the non-modified version of these aptamers were pre-incubated with rgRSV prior to infection. Lower TF was measured when rgRSV is pre-treated with modified aptamers or palivizumab, compared to infection with mock-treated RSV. The non-modified version of D4 and H9 had marginal effect on virus neutralization.

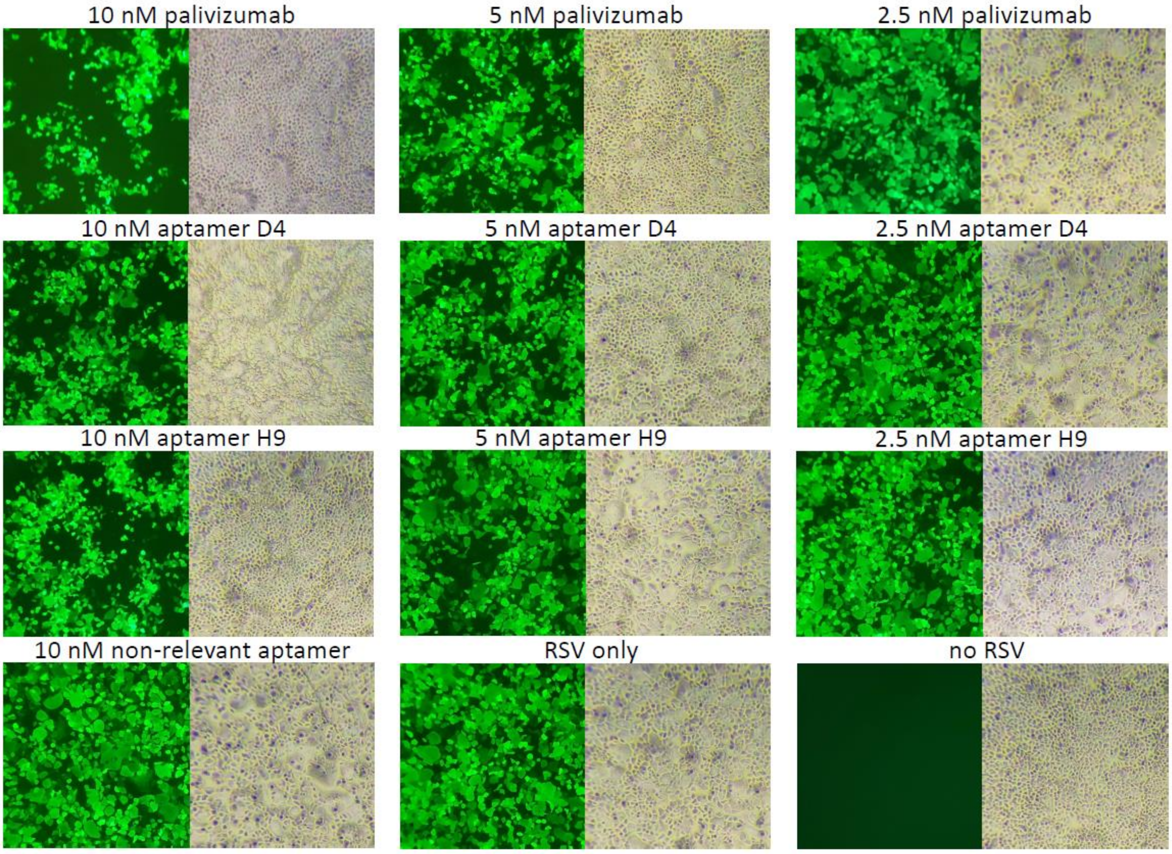


**Supplementary figure 8** Microscopy studies of rgRSV infected A549 cells 48 hrs post-infection. Green fluorescent cells and syncytia formation indicate rgRSV infection. Palivizumab, the most promising aptamers (D4, H9), and a non-relevant aptamer was pre-incubated with rgRSV prior to infection. Reduction in the total fluorescence and the number of syncytia is most visible when rgRSV was incubated with either 10 nM of palivizumab, D4 or H9.

**Supplementary table 1** $K_D$ of the 14 outstanding aptamers was determined by MST. A serial dilution was made of F proteins (prefusion F and postfusion F, labelled as pre F and post F, respectively) and mixed with a constant concentration of Cy5-labelled modified aptamers, then the IR induced thermophoresis was measured. NA means that the Affinity MO program was unable to fit a model to the obtained data points using Michaelis-Menten kinetics.

| *Aptamer* | *Target molecule* | *$K_D$ (nM)* | *± SD (nM)* |
|---|---|---|---|
| A2 | prefusion F | 123.93 | 12.90 |
| | postfusion F | 994.45 | 376.93 |
| A3 | prefusion F | 450.9 | 87.4 |
| | postfusion F | 3845.7 | 3824.7 |
| B1 | prefusion F | 1699.60 | 337.69 |
| | postfusion F | 3991.50 | 1253.30 |
| B6 | prefusion F | 11559.8 | 7767.0 |
| | postfusion F | NA | |
| B7 | prefusion F | 449.7 | 125.0 |
| | postfusion F | 1362.9 | 876.7 |
| C2 | prefusion F | 1309.5 | 433.1 |
| | postfusion F | 23405.3 | 26417.1 |
| D4 | prefusion F | 366.08 | 109.28 |
| | postfusion F | 2761.00 | 1058.90 |
| F1 | prefusion F | 863.7 | 207.4 |
| | postfusion F | 14487.5 | 32767.8 |
| F2 | prefusion F | 791.6 | 180.7 |
| | postfusion F | 59346.4 | 317154.1 |
| F5 | prefusion F | 1725.9 | 956.7 |
| | postfusion F | 10959.1 | 10913.9 |
| G6 | prefusion F | 689.59 | 272.34 |
| | postfusion F | 63662.00 | 276033.75 |
| G11 | prefusion F | 582.7 | 256.7 |
| | postfusion F | NA | |
| H8 | prefusion F | 830.79 | 149.71 |
| | postfusion F | 4062.80 | 2356.50 |
| H9 | prefusion F | 151.78 | 42.86 |
| | postfusion F | 938.06 | 487.35 |

**Supplementary table 2** Selectivity of the 6 functionally analysed aptamers was determined by MST. A serial dilution was made of viral proteins (RSV F prefusion protein (Ds-Cav1), RSV F postfusion, SARS-Cov2 spike protein (Sarscov2 S2), influenza H1 hemagglutinin (fluH1 HA), human parainfluenza virus prefusion protein (hPIV preF) and human metapneumovirus prefusion protein (MPV preF) and mixed with a constant concentration of Cy5-labelled modified aptamers, then the IR induced thermophoresis was measured. In case of most aptamers (B1, D4, G6, H8, H9) high-affinity interactions with other fusion proteins of human respiratory viruses were not detected, however, aptamer A2 interacts with SARS-Cov2 spike protein with high affinity. N.A. means that the Affinity MO program was unable to fit a model to the obtained data points using Michaelis-Menten kinetics.

| | | **RSV F selective aptamers** | | | | | |
|---|---|---|---|---|---|---|---|
| | | **A2** | **B1** | **D4** | **G6** | **H8** | **H9** |
| **Ds-Cav1 RSV prefusion** | $K_D$ *[nM]* | 123.93 | 1699.60 | 366.08 | 689.59 | 830.79 | 151.78 |
| | ±*SD [nM]* | 12.90 | 337.69 | 109.28 | 272.34 | 149.71 | 42.86 |
| **RSV F postfusion** | $K_D$ *[nM]* | 994.45 | 3991.50 | 2761.00 | 63662.00 | 4062.80 | 938.06 |
| | ±*SD [nM]* | 376.93 | 1253.30 | 1058.90 | 276033.75 | 2356.50 | 487.35 |
| **Sarscov2 S2** | $K_D$ *[nM]* | 64.60 | 25578.92 | 104778.11 | 1815.60 | 2154.20 | 1169.60 |
| | ±*SD [nM]* | 18.05 | 102830.95 | 1119857.2 | 503.65 | 1302.60 | 396.27 |
| **fluH1 HA** | $K_D$ *[nM]* | 14719.85 | N.A. | N.A. | 3576.40 | 2494.40 | N.A. |
| | ±*SD [nM]* | 17550.29 | N.A. | N.A. | 3688.80 | 3234.50 | N.A. |
| **hPIV preF** | $K_D$ *[nM]* | N.A. | N.A. | N.A. | N.A. | 3988.90 | N.A. |
| | ±*SD [nM]* | N.A. | N.A. | N.A. | N.A. | 43030.00 | N.A. |
| **MPV preF** | $K_D$ *[nM]* | 6948.60 | N.A. | N.A. | N.A. | N.A. | N.A. |
| | ±*SD [nM]* | 15655.00 | N.A. | N.A. | N.A. | N.A. | N.A. |

**Supplementary table 3** Oligonucleotide sequences

| *Name* | *Sequence (5'-3')* | *Manufacturer* |
|---|---|---|
| Library (L8) | ATC CAG AGT GAC GCA GCA 40N GAG ATA TCG TGC TAC CGT GA | IDT |
| L8 forward primer | ATC CAG AGT GAC GCA GCA | IDT |
| L8 reverse primer | TCA CGG TAG CAC GAT ATC TC | IDT |
| M13 forward | GTA AAA CGA CGG CCA G | Sigma Aldrich |
| M13 reverse | CAG GAA ACA GCT ATG AC | Sigma Aldrich |
| Non-relevant aptamer (unpublished poliovirus aptamer) | AGT GCT CCT ACT CCA CCC TAG TAA ATG TAA AAC ACC GGG TCG TGC GGG GTC TCG GCC AGG GTG AGA CTC GAT GCT G | Sigma Aldrich |
